# A moment of change: shifts in myeloarchitecture characterise adolescent development of cortical gradients

**DOI:** 10.1101/706341

**Authors:** C Paquola, RAI Bethlehem, J Seidlitz, K Wagstyl, R Romero-Garcia, KJ Whitaker, R Vos De Wael, GB Williams, NSPN Consortium, PE Vértes, DS Margulies, BC Bernhardt, ET Bullmore

**Affiliations:** Multimodal Imaging and Connectome Analysis Lab, McConnell Brain Imaging Centre, Montreal Neurological Institute and Hospital, McGill University, Montreal, QC, Canada; University of Cambridge, Department of Psychiatry, Cambridge CB2 0SZ, UK; Autism Research Centre, Department of Psychiatry, University of Cambridge, England, United Kingdom; Developmental Neurogenomics Unit, National Institute of Mental Health, Bethesda, MD 20892, USA; Frontlab, Institut du Cerveau et de la Moelle épinière, UPMC UMRS 1127, Inserm U 1127, CNRS UMR 7225, Paris, France; The Alan Turing Institute; Wolfson Brain Imaging Centre, Department of Clinical Neurosciences, University of Cambridge

**Keywords:** Adolescence, MRI, microstructure, connectivity, hierarchical gradients, transcriptomics

## Abstract

The biological processes underpinning adolescent brain maturation remain elusive. Expanding on previous work showing age-related changes in cortical morphology, we studied an accelerated longitudinal cohort of adolescents and young adults (n=223, two time points) to investigate dynamic reconfigurations in myeloarchitecture. Intracortical profiles were generated using magnetization transfer (MT) data, a myelin-sensitive magnetic resonance imaging contrast. Mixed-effect models of depth specific intracortical profiles demonstrated two separate processes i) related to overall increases in MT, and ii) showing a flattening of the MT profile related to enhanced signal in mid-to-deeper layers, especially in heteromodal and unimodal association cortices. This development was independent of morphological changes, and enhanced MT in mid-to-deeper layers was found to spatially co-localise specifically with gene expression markers of oligodendrocytes. Covariance analysis between all pairs of intracortical profiles revealed that these intracortical changes contributed to a gradual and dynamic differentiation from higher-order to lower-order systems. Depth-dependent trajectories of intracortical myeloarchitectural development contribute to the maturation of structural hierarchies in the human neocortex, providing a model for adolescent development that bridges microstructural and macroscopic scales of brain organization.

**eLife digest:** Intracortical myelin imposes a spatial structure on cortico-cortical connections, yet little is known about how myeloarchitecture develops throughout youth. We formulated a novel approach to study cortical myeloarchitecture in individual humans and leveraged an accelerated longitudinal design to track age-related changes from 14-27 years. We discovered two unique processes: one involving increasing mean myelin and another characterised by the preferential accumulation of myelin in mid-to-deeper cortical layers. Both processes contributed to an increasing segregation of lower-order from higher-order systems along the macroscale cortical hierarchy. These findings illustrate how layer specific microstructural changes contribute to the maturation of cortical organization and suggest adolescent fine tuning of hierarchical gradients of cortical networks.

## Introduction

Adolescence is a crucial phase in biological and psychosocial maturation and involves large-scale reconfigurations of brain anatomy ^1^. Previous histopathological and neuroimaging studies have shown marked age-related changes in brain structure during this sensitive period ^2–4^. This is particularly evident in magnetic resonance imaging (MRI) assessments of the macro-structural morphology of cortical regions, which have revealed regionally-variable dynamics of cortical thinning in adolescence ^2–4^. While these findings confirm the existence of strong biological forces shaping adolescent brain anatomy, morphometric analyses typically only quantify shape changes of the inner and outer cortical boundaries. In turn, these analyses may not be specific for microstructural changes occurring within the cortical mantle, which ultimately play key roles in adolescent development of cortical connectivity and function ^5^. Several recent neuroimaging studies have begun to assess intracortical microstructure in adolescence, in particular showing the utility of myelin-sensitive imaging, such as magnetisation transfer (MT), an MRI technique sensitive to the proportion of fatty tissue^6^. While recent studies confirmed associations between adolescence and changes in myelin-markers ^7–10^, findings have been mainly cross-sectional, precluding inferences on intra-individual trajectories. Furthermore, studies have generally focused on specific depths or intracortical averages, ignoring depth-dependent dynamics and thus not addressing potential systematic shifts in cortical myeloarchitecture and lamination in adolescence.

Quantitative profiling of intracortical properties across cortical depths, and specifically parameterization using central moments ^11^, has been proposed to characterise cytoarchitecture in seminal histological work ^12^ and capture inter-individual variation ^13^. In essence, studying the mean (first moment) of MT profiles perpendicular to the cortical mantle allows inferences on overall myelin content while higher order moments can address depth-dependent changes (**Figure 1**). Analysis of skewness (third moment) can contrast relative properties of deep and superficial cortical layers, and depth is a critical dimension of laminar differentiation that relates to architectural complexity ^14^ and cortical hierarchy ^15^. Applied to adolescence, such an analysis offers a non-invasive window into cortical architecture, which may recapitulate and expand classical histological findings showing overall increases in cortical myelin in adolescence ^16^ as well as transcriptional analyses suggesting that laminar signatures reflect cortical development ^17^. In addition to studying regional variations in cortical architecture, depth-dependent profiling theoretically lends a framework to tap into the large-scale coordination of different brain areas. One such approach, known as microstructure profile covariance (MPC), quantifies inter-regional similarity in microstructure at a single subject-level ^18^. Previous research has demonstrated the utility of MPC for understanding large-scale patterns of cortical architecture, specifically illustrating a sensory-fugal gradient of microstructural differentiation in both *post mortem* and *in vivo* datasets ^18^. This axis describes gradual transitions from primary sensory and motor regions with high laminar differentiation, through association cortex toward paralimbic areas with increasingly dysgranular appearance ^15^. Given prior evidence that microstructural similarity predicts cortico-cortical connectivity ^19^, tracking age-related changes in MPC provides an unprecedented way to probe coordinated maturation of microstructural networks during the critical adolescent period, moving towards a network perspective of structural brain development ^20,21^.

**Figure 1:**
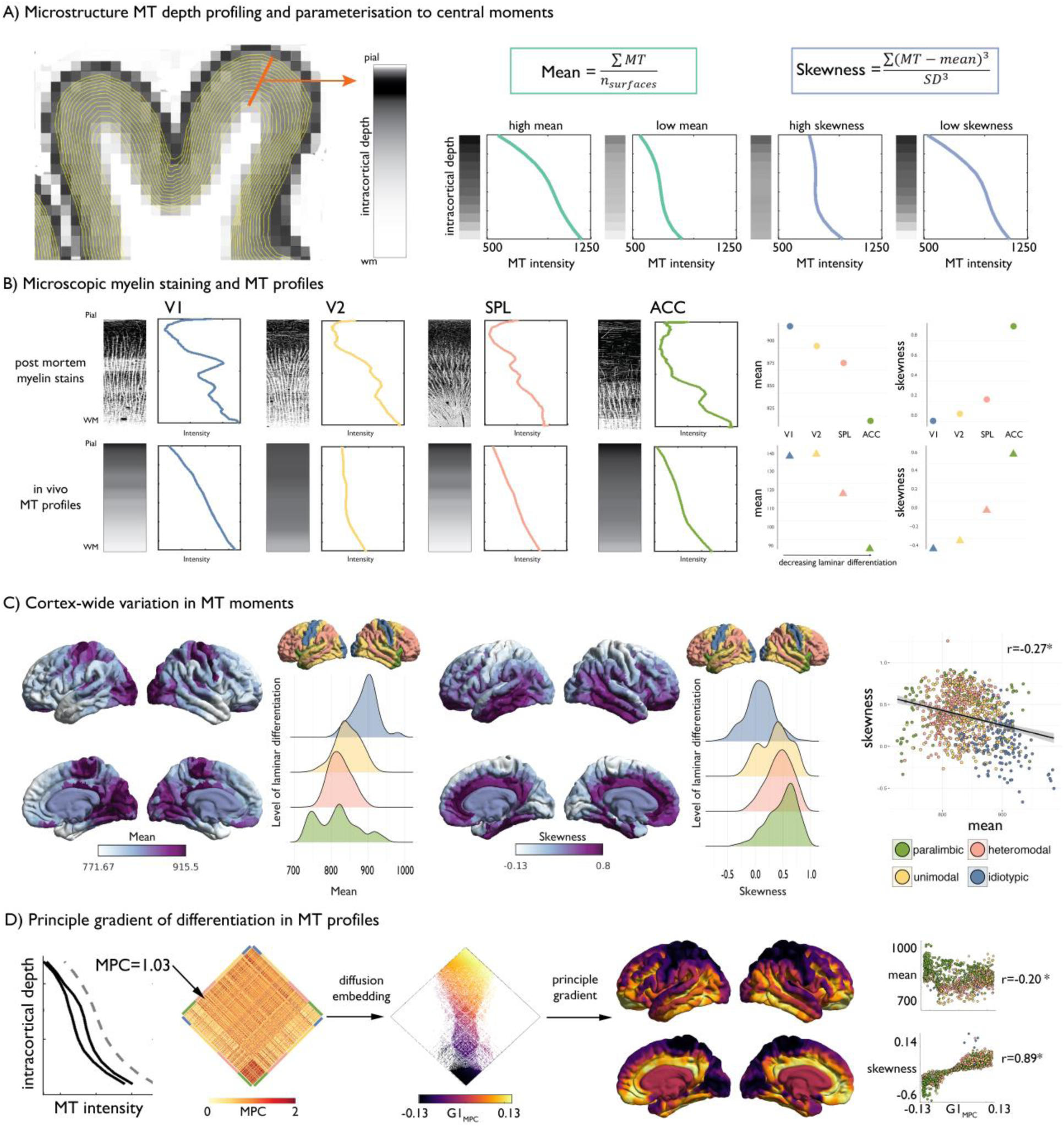
Intracortical MT depth profiling. **(A)** *Left.* Equivolumetric surfaces overlaid on an MT image of a single subject, also showing an example vertex along which MT intensity is sampled (example MT profile in grey, with lighter tones represent higher MT intensity). *Right.* Schema of first and third moments. **(B)** *Left.* Examples of microscopic myelin-stained sections (inverted image shows myelin in lighter tones) ^29,30^, with corresponding profiles and group-average MT profile from the same regions extracted *in vivo*. V1 = primary visual cortex. V2 = secondary visual area. SPL = superior parietal lobule. ACC = anterior cingulate cortex. *Right.* Dot-plots showing the moments for each exemplar profile. **(C)** Baseline group-average of first and third moment plotted on the cortical surface, and within levels of laminar differentiation^15,18^. The scatterplot shows the spatial overlap between first and third moment, coloured by level of laminar differentiation ^15,18^. Findings on the second and fourth moment are shown in **Supplementary Figures 1-2**. (D) Microstructure profile covariance (MPC) was estimated between each pair of nodes based on the partial correlation of two nodes’ MT profiles *(black)*, controlling for the cortex-wide mean intensity profile *(grey dashed)*. Baseline MPC matrices were averaged across the group, and diffusion map embedding was employed to order regions according to the principal gradient of microstructural differentiation (G1_MPC_). Scatterplots show the relationship between node-wise loadings onto the principle gradient with MT moments.

The present study examined adolescent maturation of cortical architecture and microstructure using an accelerated longitudinal NeuroScience in Psychiatry Network dataset (^22^; NSPN, 223 participants scanned twice). We translated quantitative profile analysis via statistical moment parameterisation, previously proposed for histological data, to surface-based MT imaging data and cross-referenced findings against established atlases of cytoarchitectural complexity and laminar differentiation ^23^. We tracked longitudinal changes in MT profile moments using linear mixed effect models, which leverage subject- and group-level effects to estimate microstructural changes across the entire age range ^8^. Based on histological work ^13,14,24,25^, we hypothesized that specifically the first (mean) and third (skewness) central moments of the MT profiles would capture different biological mechanisms and exhibit divergent maturational trajectories. The mean was expected to largely encompass similar changes to those captured by previous *in vivo* imaging studies on overall MT changes ^7–10^ corresponding to a shift in the overall cortical myelin content. Conversely, we anticipated skewness would capture shifts of MT intensity in the depth dimension of the intensity profile. Intracortical MT profile analysis was complemented by a series of cortical thickness and transcriptomic overlap analyses to assess morphological correlates and molecular underpinnings. In addition to studying regional MT profiles, we leveraged the MPC framework to tap into cortex-wide coordination of adolescent MT profile changes between different brain areas and consolidate our findings at a system level.

## Results

### Characterization of intracortical MT profiles

We studied the NSPN dataset, an accelerated longitudinal cohort that aggregates multimodal imaging data from 223 adolescents and young adults aged 14-27 (for details on cohort selection, processing, and quality control, see *Methods*). Briefly, we generated cortical surface models based on T1-weighted MRI and co-registered the corresponding MT volumes to these surfaces. We systematically generated equivolumetric intracortical surfaces ^26^, and sampled MT intensities along matched vertices perpendicular to the cortical mantle to build intracortical MT profiles (**Figure 1A**). Our vertex-wise technique leverages equivolumetric transformations that critically adjust surface placement in accordance to the folded cortical sheet ^26^, thereby better coinciding with the position of the putative cortical laminae than equidistant or Laplace-field guided techniques ^27^. Our intracortical approach thus offered better precision and biological validity over conventional voxel-based morphometry techniques that may be agnostic to cortical topology and intracortical architecture, and that may amplify mixing of different tissue types ^28^. MT profiles at each vertex were parcellated in 1012 approximately equal sized nodes and parameterized via central statistical moments ^14^. We focused on the first moment (mean across all cortical depths) and third moment (skewness across cortical depths), which are readily interpretable in terms of, respectively, mean myelin content and the ratio of myelination in deeper compared to more superficial layers; and have been studied in prior histological work ^14^.

### Baseline MT profiles, moments and covariance

At baseline (*i.e.*, using the average across timepoint 1), the cortex-wide average MT profile shows a non-linear increase in intensity from the superficial layers, adjacent to the pial boundary, towards the deeper layers approaching the white matter boundary, consistent with prior literature ^10^ (**Figure 1A**). The correspondence of intracortical MT profiles with myeloarchitecture was first assessed by comparing MT profiles with histological myelin stains from four regions of interest, representing the four levels of laminar differentiation ^25,29,30, 31^ (**Figure 1B, Supplementary Figure 3**, see *Methods* for details on profile quantification). Qualitatively, we observed similar variations in mean and skewness of profiles derived from *post mortem* myelin stained sections and from *in vivo* MT profiles across levels of laminar differentiation ^15^ (**Figure 1B**), supporting the extension of profile analysis from histology to *in vivo* MT imaging. At a whole cortex-level, mean MT was highest in idiotypic cortex and decreased with less laminar differentiation, while skewness exhibited an opposite pattern (spatial correlation=−0.28, p<0.001; **Figure 1C**). Negative or near-zero skewness was observed in idiotypic and unimodal areas, relating to more even distribution of MT across cortical depths, whereas more positive skewness was observed in heteromodal and paralimbic areas, related to higher MT intensities in infragranular compared to supragranular layers ^14^ (**Figure 1C**).

We explored the topology of intracortical MT patterns using microstructure profile covariance (MPC). By correlating depth-wise MT profiles, the MPC procedure estimates inter-regional microstructural similarity and has previously been validated using a combination of *post mortem* and *in vivo* data ^18^ (**Figure 1D**). Diffusion map embedding, a nonlinear dimensionality reduction technique, was employed to resolve the principle axis of microstructural differentiation. The relative positioning of nodes in this embedding space informs on similarity of their covariance patterns. In line with previous work ^18^, the first principal gradient within the baseline cohort was anchored on one end by idiotypic sensory regions and by paralimbic regions on the other end. This sensory-fugal gradient reflects systematic variations in the MT profiles, and was strongly correlated with MT profile skewness (r=0.89, p<0.001; **Figure 1D**) and weakly correlated with mean MT (r=0.20, p<0.001).

### Age-related changes in cortical depth MT profiles

Studying age-related changes in MT moments, we observed that adolescence lead to a significant increase in the mean (q_FDR_<0.00625, % rate of change: [4.8 14.7] 95% CIs; **Figure 2**), suggesting overall increases in intracortical myelin content. Conversely, we observed a unique spatial pattern of decreases in skewness (q_FDR_<0.00625, % rate of change: [−179.2 −34.6] 95% CIs; **Figure 2**). Depth-wise changes in MT intensity suggest that these decreases in skewness reflected a preferential accumulation of myelin in mid to deeper cortical layers. Controlling for overall mean MT trajectories resulted in virtually identical patterns of skewness change, suggesting relative independence between skewness and mean MT trajectories in adolescence (**Supplementary Figures 5-7**). The differential impact of age across levels of laminar differentiation was subsequently assessed by spin permutations ^21,32^. Across all moments, age-related changes in skewness were preferentially located in heteromodal (z>3.06, p<0.043) and unimodal cortex (z>3.24, p<0.024), while idiotypic nodes were less prominently represented than expected by chance (**Supplementary Figure 2 and Supplementary Table 1**).

**Figure 2:**
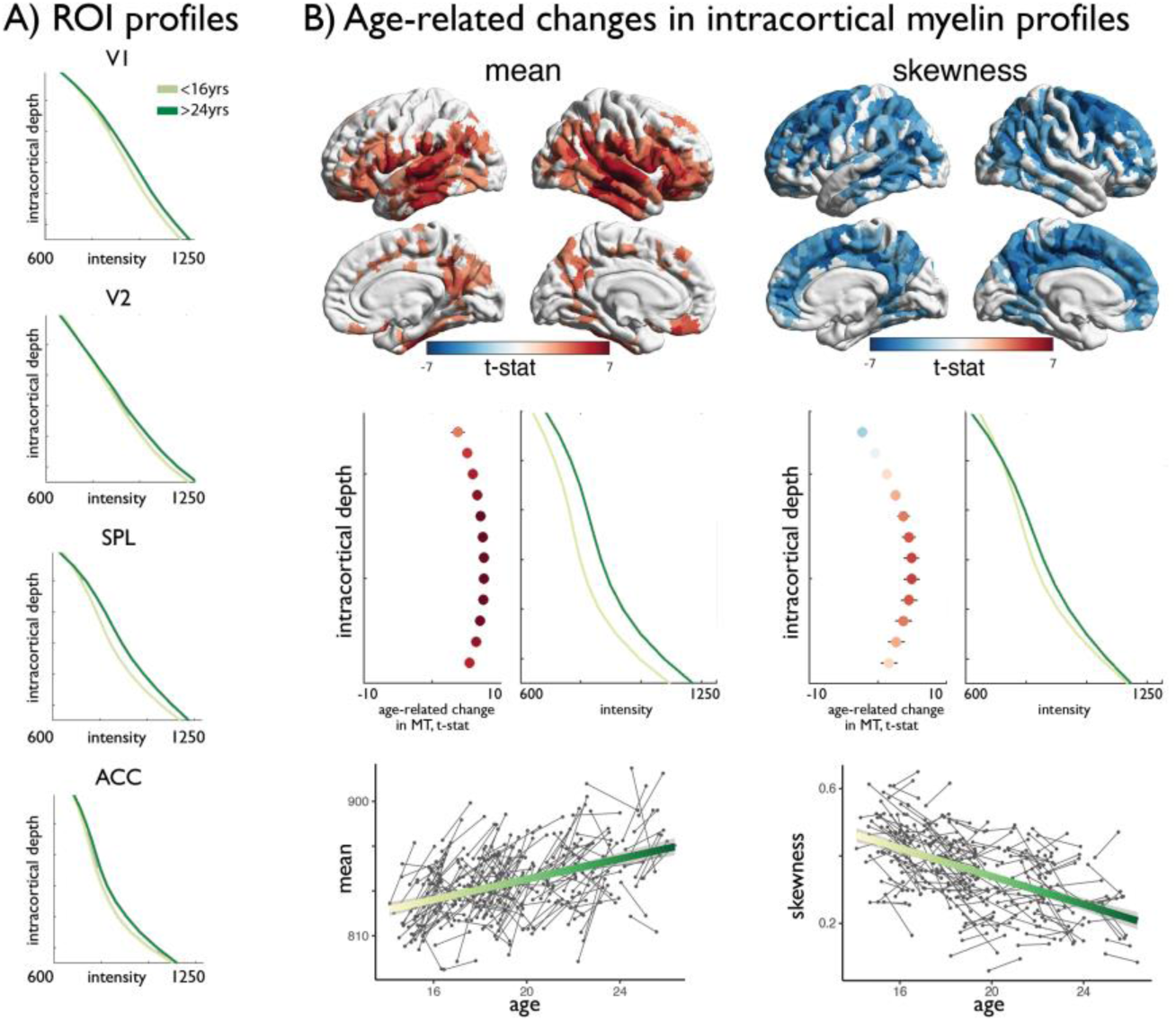
Age-related changes in MT moments. (A) Shifts in MT profiles and moments from lowest to highest age strata shown for exemplar regions of laminar differentiation (Figure 1). Moment values are provided in **Supplementary Table 3**. (B) *Upper.* Age-related changes in MT moments (qFDR<0.00625). *Middle.* t-statistic (mean ± SD) of age-related changes in MT intensity at each intracortical surface across significant regions. Mean increases were balanced across surfaces, whereas decreases in skewness were driven by preferential intensity increases at mid-to-deeper surfaces. *Lower.* Individual changes across significant regions, with regression lines depicting age-related changes across the investigated range.

### Independence of age-related intracortical MT profiles changes from cortical morphology

To examine the specificity of MT profile trajectories to intracortical variations, we assessed robustness of findings against changes in cortical morphology and boundary definition ^10,33^. To this end, we computed residual MT moments by controlling for thickness or interface blurring at each vertex within node-wise linear models. The spatial distribution of MT profile mean and skewness were virtually unchanged when data were additionally controlled for morphological and intensity confounds (r>0.99; **Supplementary Figures 5-7**). Age-related changes in MT moments were also virtually identical when using MT profile data corrected for cortical thickness and interface blurring (r>0.98, **Supplementary Figures 5-7**), supporting that MT profile trajectories were primarily driven by intracortical factors.

### Age-related change in MT profiles co-localised with expression of oligodendroglial genes

To explore molecular substrates of our imaging results, we referenced our findings against *post mortem* gene expression maps provided by the Allen Institute for Brain Sciences ^34,35^. We identified genes whose expression pattern spatially resembled the maps of the age-related change from our *in vivo* MT analysis (**Figure 2B**) ^36^. Including only those genes that passed multiple comparisons correction (p_FDR_<0.05), we conducted enrichment analysis of standard mammalian phenotypes ^37^, cell-specific expression analysis and developmental expression analysis across developmental time windows ^38^.

The pattern of age-related change in skewness showed a significant and specific transcriptomic signature (for unthresholded lists, see *Data Availability*). Cell specific expression analysis ^38^ suggested enrichment exclusively with oligodendrocytes (p=3.131e-09 [at a specificity index threshold of 0.0001]) (**Figure 3A**), confirming the association between myelination and decreased MT skewness. Developmental expression analysis ^38^ showed selective enrichment for adolescence and young adulthood (**Figure 3B**). Both analyses thus confirmed a spatio-temporal overlap of our findings from NSPN with myelin processes during adolescence. Furthermore, gene ontology analysis of genes associated with decreasing skewness was negatively associated with demyelination (Z=−2.16, p=0.0016) and axon degeneration (Z=−2.34, p=0.011), reinforcing associations to myelin. In sum, genes negatively associated with age-related skewness reduction were less likely to be enriched for demyelination and were exclusively associated with oligodendrocytes. Given that the present data is based on healthy cortex, those same areas are not high in expression of “demyelinating” genes, but instead show a negative signal (*e.g.*, negative z-scores). This is expected since those areas of high expressing myelin genes are most vulnerable ^32^. Thus, MT skewness changes during adolescence development appear to be strongly reflective of changes in myelination. None of the other moments provided a list of genes passing the threshold set by the false discovery rate procedure. However, exploratory rank based correlations on the weights of each gene indicated a small correlation between mean and skewness related genes (rho=0.14, p<10e-16), indicating mean and skewness share a modest transcriptomic signature. Finally, it should be noted that the same analysis on baseline data only (average across a single scan per subject) from this cohort did not identify a whole genome transcriptional profile that was co-located with any of the architectonic moments of cortical micro-structure (**Supplementary Figure 8**), reinforcing the added value of developmental data (e.g. measuring an age-related change) in detecting human neurodevelopmental processes by myelin-sensitive MRI.

**Figure 3:**
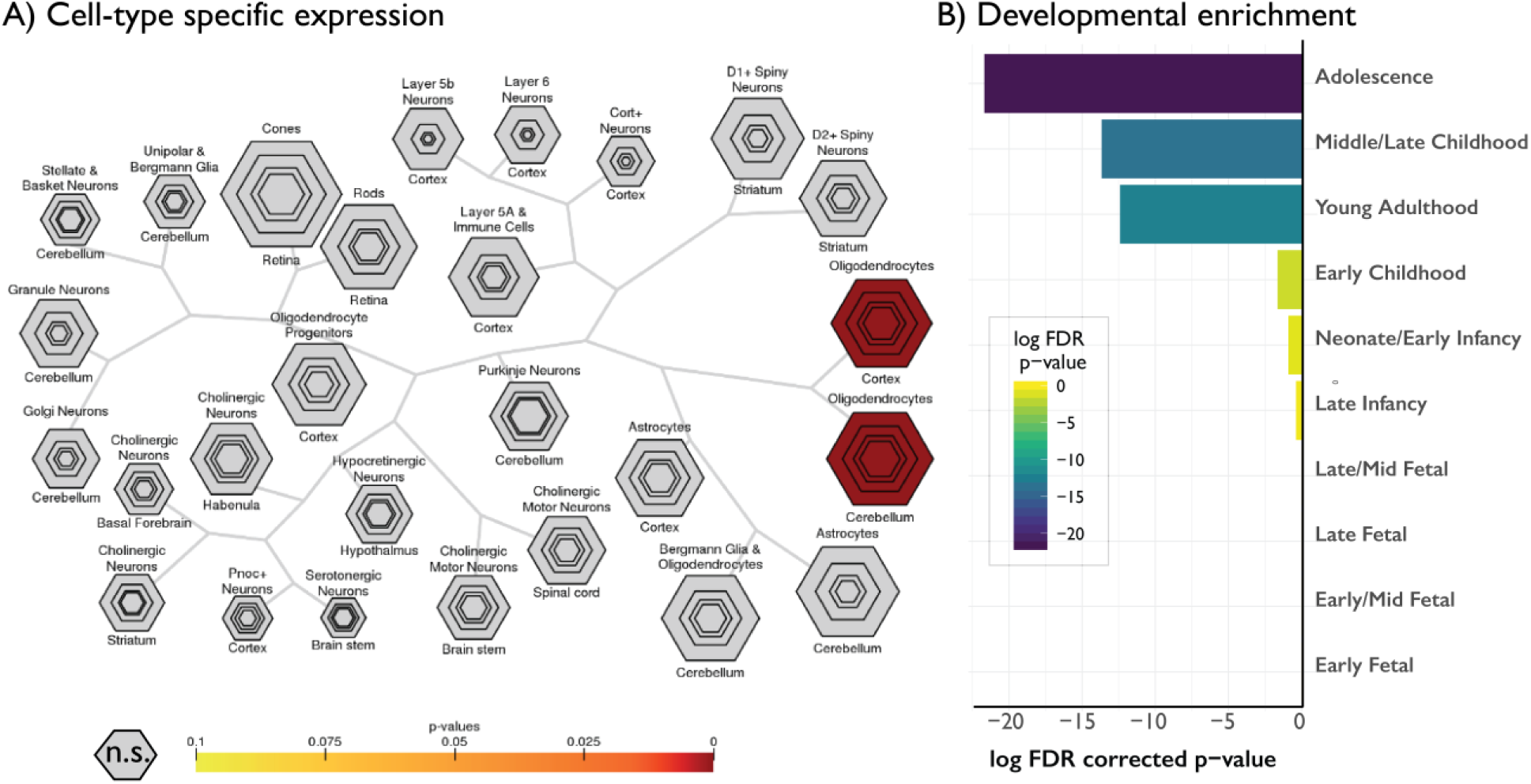
Gene decoding of age-related changes in skewness against the Allen Institute for Brain Sciences gene expression atlas. Only genes negatively relating to age-related changes in skewness are shown as only these genes survived FDR < 0.05. None of the other four moments had any significantly associated genes. Cell-type specific expression, showing selective enrichment for oligodendrocytes, reinforcing the link between profile skewness and myelination. Hexagon rings denote significance at different pSI thresholds (from p < 0.05 in the outer ring to 0.00001 in the center). **B)** Developmental cortical enrichment, showing enrichment specifically in adolescence and young adulthood.

### Adolescent reconfiguration of microstructure profile covariance networks

Adopting the MPC framework, we subsequently assessed inter-regional coordination of myeloarchitectural development. MPC networks were constructed for each participant and we modelled age-related changes in microstructural similarity between all node pairs (**Figure 4A**). At the node-to-node edge-level, adolescent development was primarily related to increases in MPC (1564 increases vs 534 decreases at q_FDR_<0.025) (**Supplementary Figure 9A**). To express the spatial pattern of myeloarchitectural development in a lower dimensional, and more readily interpretable space, we implemented diffusion map embedding ^39^ (**Figure 4A**). Nodes closer in this embedding space increase in microstructural similarity during adolescence, whereas distant nodes decouple. The first principal component (G1_DEV_) illustrated a sensory-fugal gradient, explaining 33% variance (**Figure 4A**). On one end of the gradient, idiotypic sensory and motor areas became increasingly coupled in adolescence, and more segregated from the opposite anchor constituted mainly of paralimbic nodes. The concordance of the developmental gradient (G1_DEV_) with the baseline MPC gradient (r=0.89, p<0.001, **Figure 1D**) suggested that the axis of microstructural differentiation expands during adolescence. To synoptically visualise these dimensional changes, we generated, aligned, and contrasted cross-sectional MPC gradients (G1_MPC_) within the youngest (<16 years) and oldest (>24 years) age strata. Similar results were observed using alternative age windows. Heteromodal and unimodal cortex, occupying central regions of the gradient, were drawn outwards towards one of the gradient anchors in older individuals (**Figure 4B**), forming a more bimodal distribution (**Supplementary Figure 10**). Specifically, prefrontal and medial parietal areas were increasingly coupled with sensory areas, and temporal regions increased in similarity to paralimbic regions (**Figure 4B**). By relating the age-related shifts in MPC to anchors with age-related changes in MT moments, we found that greater decreases in MT skewness were related to sensory anchor coupling (r=0.57, p<0.001; **Figure 4C)**. Conversely, restricted age-related skewness changes were associated with paralimbic anchor coupling (r=0.62, p<0.001). Together, these findings demonstrate how depth-dependent myelination during adolescence reshapes myeloarchitecture and underpins macro-scale reorganisation of cortical networks.

**Figure 4:**
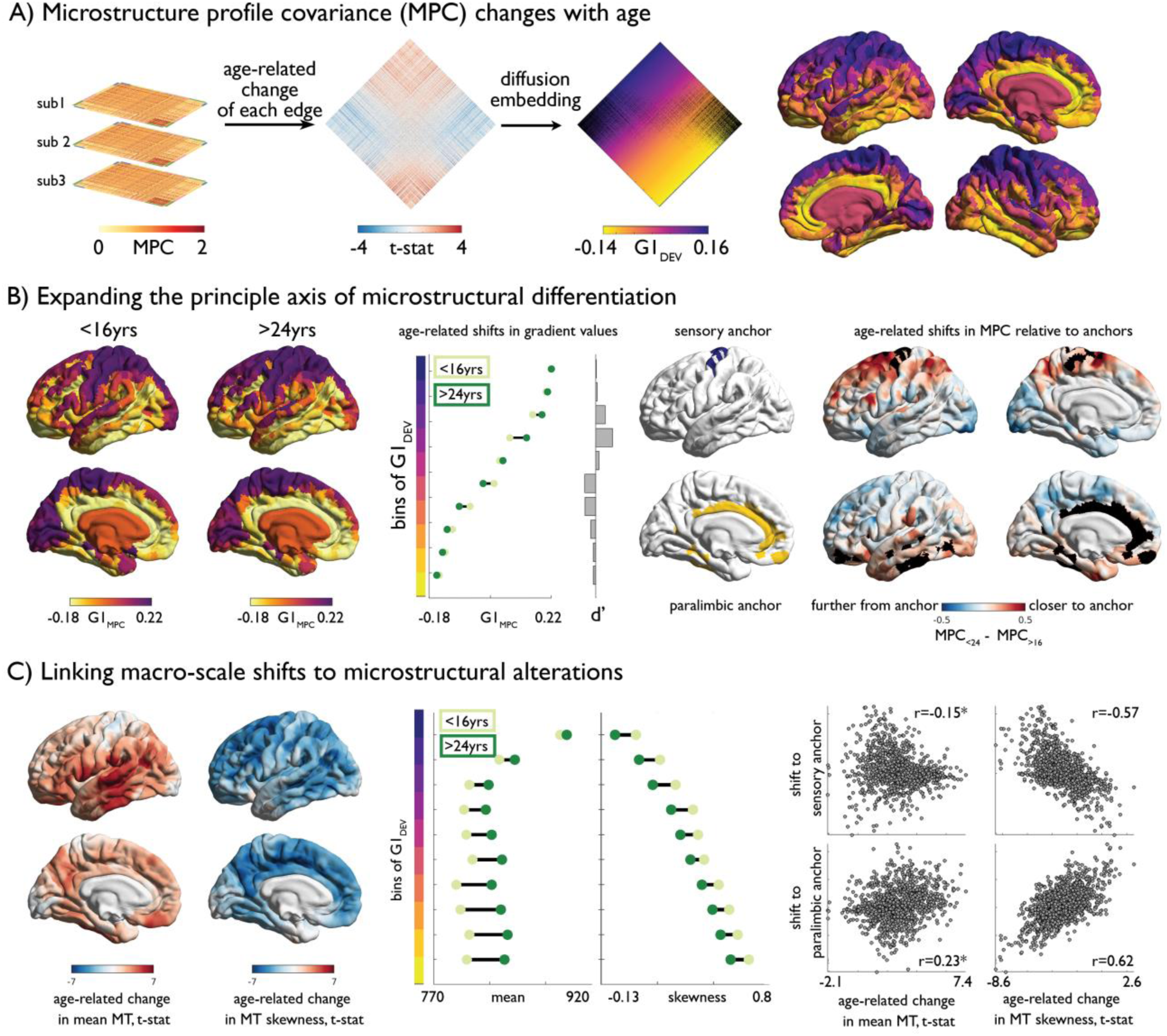
Age-related changes in microstructure profile covariance (MPC). **(A)** Subject-specific MPC matrices (*stacked)* were used in mixed effect models to calculate age-related changes in microstructural similarity between each node pair, generating a t-statistic matrix *(middle)*. Diffusion map embedding ordered this matrix along the principal axis (G1_DEV_) *(right)*. Rows of the matrix were coloured according to G1_DEV_. Surface projection of G1_DEV_ illustrates a transition from primary sensory *(purple)* through association *(pink)* to limbic cortices *(yellow)*. **(B)** *Left.* Principal axis of microstructural differentiation within extreme age strata (G1_MPC_), show a sensory-fugal axis similar to the baseline and developmental analysis. *Middle.* Shifts in average G1_MPC_ within ten discrete bins of G1_DEV_, and corresponding Cohen’s d effect size. Central regions of G1_DEV_, aligning with association cortices, expand from the centre of G1_MPC_, towards either sensory or paralimbic anchors. *Right.* Age-related shifts towards anchors visualised via age-strata difference in MPC. The average MPC to each anchor was calculated within each extreme age-strata then subtracted. Prefrontal and parietal areas increased in microstructural similarity with the sensory anchor (*top*), whereas temporal regions became more coupled with the paralimbic anchor. **(C)** *Left.* Age-related change in MT moments (unthresholded maps from Figure 2B). *Middle.* Age-related shifts in mean and skewness within each bin of the gradient. *Right.* Correlation between age-related change in MT moments (Figure 2B) and shifts to anchors (Figure 4B right).

## Discussion

Our longitudinal analyses revealed marked alterations in intracortical myeloarchitecture during adolescence, which was accompanied by large-scale reorganisation of microstructural gradients. Changes to intracortical myelin can be generally characterised by two developmental processes, one involving overall increases in mean MT and one involving preferential MT increases in mid-to-deeper cortical layers. Both processes particularly affected unimodal and heteromodal association cortices *i.e.*, areas between low-level sensory and motor areas on the one end of the cortical hierarchy and high-level transmodal and limbic regions on the other end. A series of complementary explorations, based on alternative neuroimaging features known to undergo large-scale changes in adolescence ^2–4^, demonstrated the independence of these intracortical microstructural changes from age-related effects on overall cortical morphology and tissue contrast. In addition, cross-referencing our findings against *post mortem* gene expression maps showed that patterns of age-related change significantly and specifically overlapped with genes associated with oligodendrocyte processes, supporting the specificity of MT changes to myelin. Leveraging microstructure profile covariance analysis, we explored coordination of intracortical microstructure characteristics across distributed areas ^18^. In doing so, we showed that local processes were paralleled by a cortex-wide differentiation along the sensory-fugal axis. This axis ultimately underpins the segregation of lower-from higher-order components of the putative cortical hierarchy.

Inspired by classical histological studies operating on 2D sections ^12–14^, our work formulated a surface-based approach to quantify myeloarchitecture *in vivo* via systematic profiling of intracortical MT intensities in the direction of cortical columns and its parameterization via the profiles’ central moments. The skewness of the MT profile, reflecting nonlinear increases in myelin towards deeper layers ^27,40,41^, tilts from infragranular-dominant in paralimbic regions towards a more even distribution across cortical depths in sensory regions. This graded shift parallels changes in the laminar origin of projections, where paralimbic regions engaged in feedback processing project from infragranular layers and sensory regions providing feedforward information project predominantly from supragranular layers ^42^. Meanwhile, mean myelin increased from polar towards sensory regions, recapitulating prior *post mortem* evidence ^29,30^, established atlases of cytoarchitecture ^23^, and resting-state functional connectivity gradients ^39,43^. These baseline analyses demonstrate the power of this framework to profile human myeloarchitecture *in vivo*.

Tracking longitudinal change in intracortical MT profiles, we found concomitant but independent developmental patterns in the developmental trajectory of mean and skewness. While both sets of age-related changes were strongly and preferentially located in heteromodal and unimodal association cortex, they only partially overlapped which was also supported by gene expression analysis. Indeed, only skewness but not mean myelin changes were driven by oligodendrocyte modulation ^24^, and these effects were found to be independent from interface blurring and cortical thickness. A recent review ^24^ proposed mechanisms of experience-dependent myelination, which appear to align with our findings. Accordingly, increasing mean likely reflects myelination of previously unmyelinated axons. On the other hand, compositional changes in myelin architecture of axons would not lead to overall in- or decreases in myelin signal per se. Given skewness changes were independent from mean changes, these more likely capture architectural reconfigurations. Underlying mechanisms may include *de novo* generation of oligodendrocytes ^45^, activity-dependent modifications in neuronal and oligodenodrocyte precursors ^46^, and tuning in the distance between nodes of Ranvier along both cortico-cortical as well as white matter axons ^47^. Building on experimental evidence in animals suggesting that experience-induced oligodendrocyte remodeling occurring during this time may be critical for overall brain function and mental health ^48^, and human imaging findings showing perturbations in myeloarchitecture across several brain disorders ^33,49,50^, these architectural changes may thus represent a microstructural context that could determine susceptibility to a range of neuropsychiatric and neurological conditions occurring in late childhood and adolescence.

Adaptation of a recent microstructure profile covariance analysis framework to longitudinal MT data illustrated the macroscale impact of microstructural changes. Specifically, decreases in MT profile skewness pushed regions to a more sensory-like architecture (which was coupled with high mean MT). In contrast, increases in skewness pushed regions towards a paralimbic-like MT profile. These regions also underwent protracted increases in mean MT, but still did not reach the high levels observed in sensory regions. These two regionally distinct developmental patterns within the association cortex promoted a more bimodal distribution of myeloarchitectural types across the cortex. Similar properties of modular segregation in tractography-based networks have been shown to mediate age-related improvements in executive function throughout adolescence ^51^, reinforcing the importance of these processes for understanding psychosocial maturation. Drifts in association cortex towards either sensory-like or paralimbic-like architecture represents, thus, an expansion of the sensory-fugal gradient of microstructural differentiation ^18^. Such a sensory-fugal gradient was previously described by Mesulam ^15^ based on non-human primate research to encapsulate cortex-wide variations in architecture and connectivity, and has since been suggested to reflect increasing synaptic plasticity towards transmodal regions ^52^. Systematic variations in the degree of experience-dependent plasticity may explain why myelin-derived markers develop differently along the sensory-fugal gradient ^24^, in contrast to other known developmental gradients such as the rostro-caudal timing of terminal neurogenesis ^53^. Differentiation of association cortices also conforms with notions of the “tethering hypothesis” of cortical evolution ^54^. According to this framework, intermediate regions of the cortical hierarchy are less constrained by extrinsic inputs and intrinsic signaling molecules, which has allowed for massive surface area expansion throughout mammalian evolution. Our findings suggest that reduced constraints on association cortices allow for protracted development of myeloarchitecture, whereas the sensory and limbic anchors are well-defined prior to adolescence.

Our work suggests that myelination during adolescence is unlikely to be a question of simply more or less. Instead, our longitudinal findings show that the type of change is topologically divergent when we take depth into consideration. Expanding upon previous evidence of adolescent increases in overall mean intracortical myelin content, our findings demonstrate a relative preferential specific accumulation of myelin towards mid-to-deeper infragranular layers mainly in association cortices. As our analyses have shown, these findings are not explainable by non-specific changes in overall cortical morphology during adolescence, but instead likely reflect architectural changes associated with oligodendrocyte related processes. The coordinated change patterns strengthen the notion that the forces of adolescent development further widen the existing axis of macroscale cortical organization, driving association cortex either closer towards sensory or limbic systems. Thus, our findings illustrate how cell-type and layer specific microstructural changes assessed in the direction of cortical columns contribute to the maturation of macroscale cortical organisation and suggest adolescent calibration of structural hierarchical gradients.

## Methods

### Imaging acquisition and processing

Details on this cohort and scanning have been described in detail elsewhere ^10^. Briefly, T1w and MT data were acquired from 223 healthy adolescents at two time points as part of the NeuroScience in Psychiatry Network’s accelerated longitudinal study (baseline age range = 14-25, inter-scan interval = +/-18 months, MPM sequence ^55^ on three identical TIM Trio 3T scanners; two located in Cambridge and one located in London). The MPM sequence comprises several multi-echo 3D FLASH (fast low angle shot) scans ^55^. MT-weighting was achieved by applying an off-resonance Gaussian-shaped RF pulse (4 ms duration, 220° nominal flip angle, 2 kHz frequency offset from water resonance) prior to the excitation with TR/α = 23.7 ms/6°. Multiple gradient echoes were acquired with alternating readout polarity at six equidistant echo times (TE) between 2.2 and 14.7 ms. for MT weighted acquisition. Other acquisition parameters were: 1 mm isotropic resolution, 176 sagittal partitions, field of view (FOV) = 256 × 240 mm, matrix = 256 × 240 ×176, parallel imaging using GRAPPA factor 2 in phase-encoding (PE) direction (AP), 6/8 partial Fourier in partition direction, non-selective RF excitation, readout bandwidth BW = 425 Hz/pixel, RF spoiling phase increment = 50°.

FreeSurfer (v5.3) pre-processing was used to reconstruct pial and white matter surfaces for each participant based on the T1w scans and all subjects underwent extensive quality control with manual editing where possible (see ^10^ for details). Following cortical surface reconstruction and surface-based co-registration between T1w and MT weighted scans, we generated 14 equivolumetric cortical surfaces within the cortex ^26^, and systematically sampled MT intensity along these surfaces ^18^ (**Figure 1**). Next, depth-wise MT profiles were calculated across all vertices in native space and MT profiles were averaged within 1012 equally sized nodes (**Figure 1**). The parcellation scheme was mapped from a standard space (fsaverage) to native space for each subject using surface-based registration ^43^). We corrected for depth-specific partial volume effects (PVE) of cerebrospinal fluid using a mixed tissue class model ^56^ to reduce potential bias of averaging MT values in voxels with CSF ^57^. Specifically, we fitted a linear model at each node (n) and each surface (s) of the form

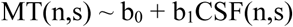

where MT(n,s) and CSF(n,s) represent node-specific, surface-specific MT value and CSF partial volume effect estimates. Final CSF-corrected MT values were calculated as the sum of the residuals [MTc(n,s) = T1(n,s) – (b0+b1*CSF(n,s))] and the uncorrected group average MT value.

### Baseline analysis of MT profiles

We characterised the MT profiles by the central moments of the intensity distributions; mean, standard deviation (SD), skewness, and kurtosis ^14^. We focused our main analysis on the first and third moments (**Figure 1**), owing to the collinearity of SD with mean and kurtosis with skewness (see also **Supplementary Figure 4**), the clearer biological interpretation of mean and skewness for the MT profiles, as well as how the first and third moments capture different dimensions of the intensity distribution. Prior to statistical assessment, we identified and removed outlier individuals (n=33). Outliers were defined as individuals with >1% nodes that deviate from the age-strata median of mean MT by >3 interquartile ranges. In an effort to validate the relationship between MT and myelin, which is discussed in more detail elsewhere ^6,55,58^, we examined MT-derived and *ex vivo* derived myelin profiles in matched regions of the cortex. We selected hyper-stained myelin pictures from classical literature ^29,30^ that represent different levels of cytoarchitectural complexity and have been well characterised in recent work ^25^. We extracted a rectangle section of each image (spanning from the pial layer to the white matter boundary; **Figure 1A**), inverted the colour to align with MT imaging (*i.e.*,: myelin is more white), obtained intensity values per pixel by reading the image into MATLAB (v2017b), then generated a region-specific intensity profile by averaging values row-wise. Finally, we calculated the MT moments for each region and contrasted these values with baseline group-average MT moments for matched regions. Additionally, we contrasted baseline group-average MT moments across levels of laminar differentiation ^18,31^ and cytoarchitectonic type ^10,23^ to determine whether the *in vivo* derived MT moments recapitulate histological evidence of cytoarchitectonic variation. MPC networks were calculated as the pairwise Pearson correlation between nodal MT profiles, controlling for the average whole-cortex MT profile.

### Longitudinal assessment of age-related changes in MT moments

Age-related changes in MT moments were estimated at each node within four linear mixed effect models (LME) using SurfStat ^59^ for Matlab, accounting for the non-independence of subjects as well as sex, using the following model:

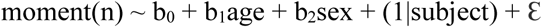

where *n* represents the node. The addition of person-specific random intercepts significantly improves model fit in accelerated longitudinal designs. LMEs were performed on each node and t-statistics were projected onto the cortical surface. Additionally, we examined the relationship between level of laminar differentiation and adolescent development of MT moments. To assess the relative over- or under-representation of significant regions per laminar class we generated 10,000 spin permutations of the *t*-maps, which controls for spatial contiguity and hemispheric symmetry across permutations ^21,32,44^. Specifically, we differently rotated the *t*-maps 10,000 times and for each permutation of this spin or rotation we computed counts per laminar class that passed a false discovery rate (FDR) correction for multiple comparisons ^62^ to create a null-distribution of the counts table. Based on the null distribution, we computed *Z*-scores and two-sided *p*-values for the actual count table and corrected for multiple comparisons across the entire table using conservative Bonferroni correction.

### Gene expression analyses

We measured the spatial overlap of our imaging derived baseline (*e.g.*, one scan per subjevy) MT moments (**Figure 1E**) and t-statistic maps (**Figure 2A**) with *post mortem* Allen Institute for Brain Sciences (AIBS) gene expression maps ^36^. We used Bayesian random effects analysis of AIBS gene expression with spatial topology, as implemented in NeuroSynth gene decoding ^34,35^, to obtain a list of genes significantly associated with each of the *t*-maps ^36^. We only included genes that passed FDR correction (*p*_FDR_ < 0.05) and conducted enrichment analysis of standard mammalian phenotypes (*i.e.*, observable morphological, physiological, behavioral, and other characteristics of mammalian organisms that are manifested through development and lifespan) in the Mouse-Genome Informatics database ^37^. Enrichment analyses were conducted using Enrichr ^60,61^, using a *Z*-score modification of Fisher’s exact test and FDR correction.

### Spatial topography of age-related changes in MT profiles

Age-related changes in inter-regional microstructural similarity were assessed by applying the same LME to individual edges of subject specific MPC networks ^18^. Diffusion map embedding, a non-linear dimensionality reduction technique ^63^, was applied to the resultant t-statistic matrix to discern the spatial topography of age-related changes. The first principal component represents the principal axis of variation in age-related changes in MPC. Nodal loadings onto the principal component, otherwise known as gradient values (G1_DEV_), depict the similarity of nodal patterns of MPC change. Similar to previous analyses ^18^, we examined whether G1_DEV_ differed across levels of laminar differentiation using ridge plots ^64^. For closer inspection of MT profile changes across the developmental gradient, we generated group-average MT profiles for the youngest (<16years, n=43) and oldest (>25years, n=30) age strata. To probe the impact of developmental differentiation on the maturity of the underlying microstructure map, we generated group-average MPC matrices within the youngest, oldest as well as a mid-range (20-22 years) age-strata. MPC matrices were subjected to diffusion map embedding, then the young and old embeddings were aligned to the mid-range embedding using Procrustes rotation ^65^. Next, we rank ordered and binned G1_DEV_ into ten, approximately equal sized bins (nodes per bin∼110). Age-related differences in MPC gradients were assessed by (i) bin-wise difference average G1_MPC_ values, (ii) bin-wise Cohen’s d effect size change of gradient values, (iii) node-wise difference in MPC to the gradient anchors, that is the extreme bins, and (iv) bin-wise difference in MT moment values. To reconcile macroscale changes in the MPC gradients to microstructural alterations, we performed a spatial Pearson correlation between unthresholded t-statistic maps of age-related changes in MT mean and skewness (from Figure 2B) with shifts relative to anchors (map iii, Figure 4B left).

## Data and code availability

Preprocessed microstructure profiles are available via GitHub: https://github.com/MICA-MNI/micaopen/tree/master/a_moment_of_change

The repository also includes Matlab and R scripts to reproduce the primary analyses and figures.

## Acknowledgements

CP is supported by the Transforming Autism Care Consortium (TACC) and the Fonds de la Recherche du Quebec – Santé (FRQS), RAIB is supported by a British Academy Postdoctoral fellowship and Autism Research Trust. CP, RAIB, GW, BB are supported by a Cambridge-MNI collaborative research grant. RRG was funded by the NSPN and the Guarantors of Brain. JS was supported by the NIH Oxford-Cambridge Scholars’ Program. KW was supported by the Health Brain Healthy Lives (HBHL) Initiative. KJW was funded by a The Alan Turing Institute under the EPSRC grant EP/N510129/1. PEV was supported by the MRC (MR/K020706/1), is a Fellow of MQ: Transforming Mental Health (MQF17/24) and is a Fellow of the Alan Turing Institute funded under EPSRC (EP/N510129/1). BCB is supported by National Science and Engineering Research Council of Canada (NSERC, Discovery-1304413), the Canadian Institutes of Health Research (CIHR, FDN-154298), the Azrieli Center for Autism Research of the Montreal Neurological Institute (ACAR), SickKids Foundation (NI17-039), and received salary support from FRQS (Chercheur Boursier Junior 1). EB is supported by a NIHR Senior Investigator award.

Data were curated and analysed using a computational facility funded by an MRC research infrastructure award (MR/M009041/1) and supported by the NIHR Cambridge BRC. This work was supported by the Neuroscience in Psychiatry Network (NSPN) Consortium, a strategic award from the Wellcome Trust to the University of Cambridge and University College London (095844/Z/11/Z); by the Cambridge NIHR Biomedical Research Centre.

The views expressed are those of the authors and not necessarily those of the NHS, the NIHR or the Department of Health and Social Care.

## Notes

#### Summary of Updates

Spelling mistake in author list

