## Supplement for "A moment of change: shifts in myeloarchitecture characterise adolescent development of cortical gradients" (from another version of this preprint: version 1, 2019-07-23)

### Supplementary materials


**Baseline properties of SD and Kurtosis** 1

Supplementary Figure 1: 1

**Age-related changes in SD and Kurtosis** 1

Supplementary Figure 2: 2

**Cytoarchitectural complexity replication** 2

Supplementary Figure 3: 3

Supplementary Table 1: 4

Supplementary Table 2: 4

Supplementary Table 3: 4

**Correspondence between MT moments** 5

Supplementary Figure 4: 6

**Independence of age-related changes from cortical morphology** 7

Cortical thickness 7

Supplementary Figure 5: 7

Interface blurring 8

Supplementary Figure 6: 8

Global mean shifts 8

Supplementary Figure 7: 8

**Baseline gene ontology** 8

Supplementary Figure 8: 9

**Deconstructing age-related changes in MPC by cytoarchitectural class** 10

Supplementary Figure 9: 10

**Increasing bimodality of the principle gradient of microstructural differentiation** 11

Supplementary Figure 10: 11

**Developmental gradients derived from moment difference** 12

Supplementary Figure 11: 12

**References** 12

### Baseline properties of SD and kurtosis

Analogous to our main analysis on mean and skewness of myelin profiles we also assessed the properties of the other two central moments of the intensity distribution (SD and Kurtosis). Broadly we find that the SD captures a very similar pattern to the mean (spatial overlap: mean~SD: $r=0.62$, $p<0.001$), whereas kurtosis was correlated to skewness ($r=0.58$, $p<0.001$). Unsurprisingly, both moments also showed similar distributions across classes of laminar differentiation.

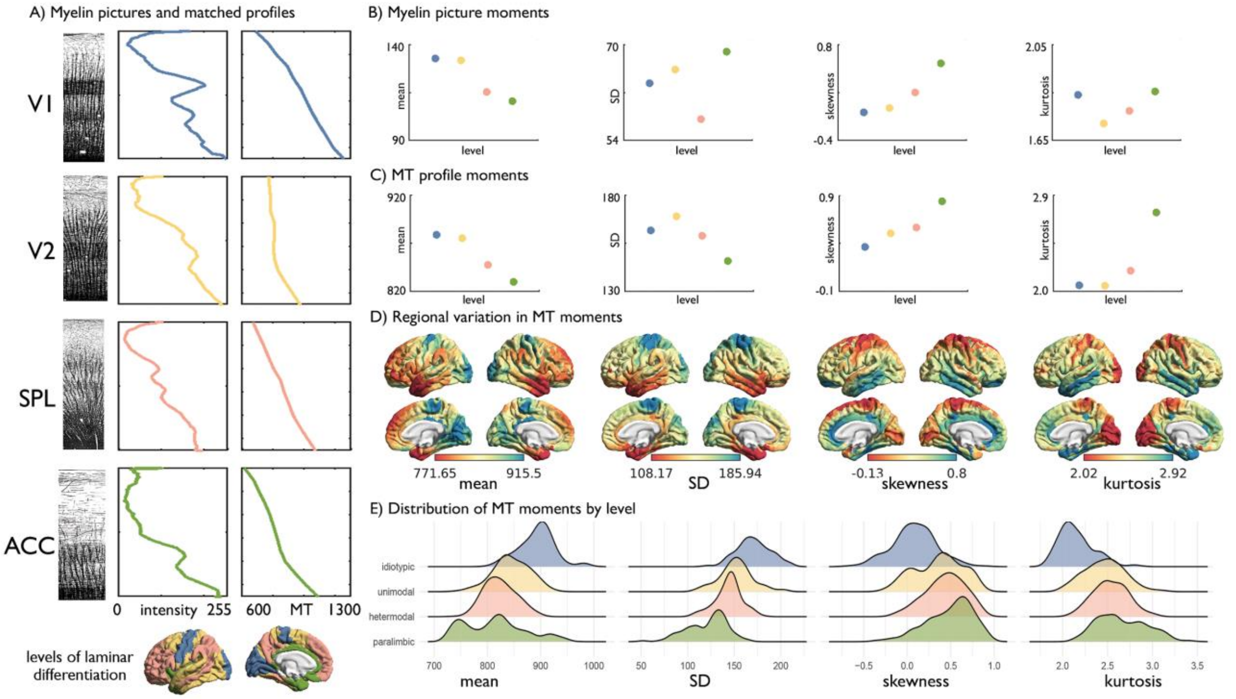


### Supplementary Figure 1:

Characterisation of intracortical MT profiles. (A) Examples of hyper-stained myelin pictures [1–3] with corresponding myelin profile and group-average MT profile from the same region. V1 = primary visual cortex. V2 = secondary visual area. SPL = superior parietal lobule. ACC = anterior cingulate cortex. Atlas of laminar differentiation projected on the cortical surface [1,2]. (B) Moment values across levels of laminar differentiation from the myelin pictures shown in (A). (C) Moment values from across levels of laminar differentiation from the MT profile shown in (A). (D) Group-average MT moment per node projected onto the cortical surface. (E) Density plots depict the distribution of group-average MT moments for each node within levels of laminar differentiation.

### Age-related changes in SD and kurtosis

Assessing age-related changes across all four moments revealed a similar pairing between the mean and SD on the one hand and skewness and kurtosis on the other. SD showed a steady increase during adolescence and preferentially impacted heteromodal cortices (z=3.42, p=0.012), while paralimbic cortex was under-represented (z=-5.60, p<0.001) as assessed using spin premutation and count tables of significant regions per class of laminar differentiation. Kurtosis showed a steady and robust decrease during adolescent development that preferentially affected heteromodal cortex (z=8.80, p<0.001). At a whole-brain level there was a moderate but significant overlap in the t-maps of age-related changes (**Supplementary Figure 4**).

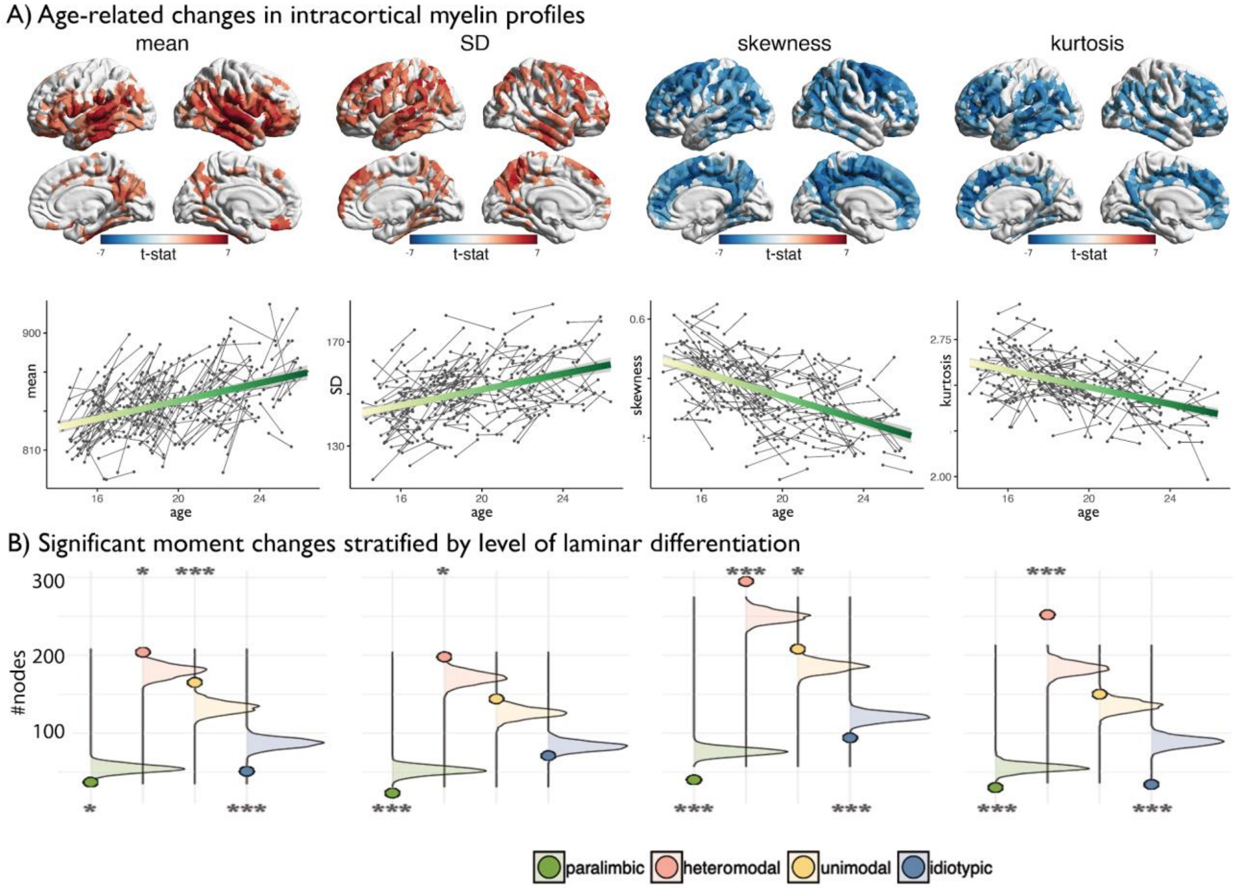


**Supplementary Figure 2:**

Age-related changes of MT moments; mean, standard deviation (SD), skewness and kurtosis. (A) Upper *t*-statistics projected on the cortical surfaces depict significant age-related changes in MT moments. Lower Scatter plots depict individual changes in average moment value across significant regions, with regression lines depicting the age-related changes. (B) Bar plots show the z-scores of the number of regions passing FDR correction for each class relative to a null-distribution generated from spin permutations of the t-statistic map (see Methods for details). Significance is indicated by asterixis and based on Bonferroni corrected p-values across all moments and classes (*** = p < 0.001, * = p < 0.05). Positive significant z-scores indicate that the number of regions passing FDR for that class are significantly higher than expected by chance, negative significant z-scores indicate the number is significantly lower (tables provided below).

**Cytoarchitectural complexity replication**

To show that neither the distribution of baseline moments (Figure 1E) nor age-related effects (Figure 2) were unique to the parcellation scheme or cortical classes [4], we replicated our findings using the Von Economo atlas of cytoarchitectural complexity [5]. We stratified the group-mean maps for each moment by Von Economo structural type [5] and again observed that the ordering of classes shows opposite patterns for the mean and SD on one hand and skewness and kurtosis on the other hand (**Supplementary Figure 3A**). Additionally, the ordering of classes from motor and sensory at one end to insula and limbic at the other end supports a sensory-fugal differentiation of MT moments. To assess the relative over- or under-representation of significant regions per structural type in the Von Economo atlas we used the spin permuted [6–8] t-maps and computed counts per structural type that passed FDR correction to create a null-distribution of the counts table. Based on the null distribution, we computed z-scores and two-sided p-values for the actual count table and corrected for multiple comparisons across the entire table using Bonferroni correction (**Supplementary Figure 3B, Supplementary Table 2**). Here, we find that age-related changes in skewness are over-expressed in association cortices and strongly under-expressed in primary and secondary sensory cortices.

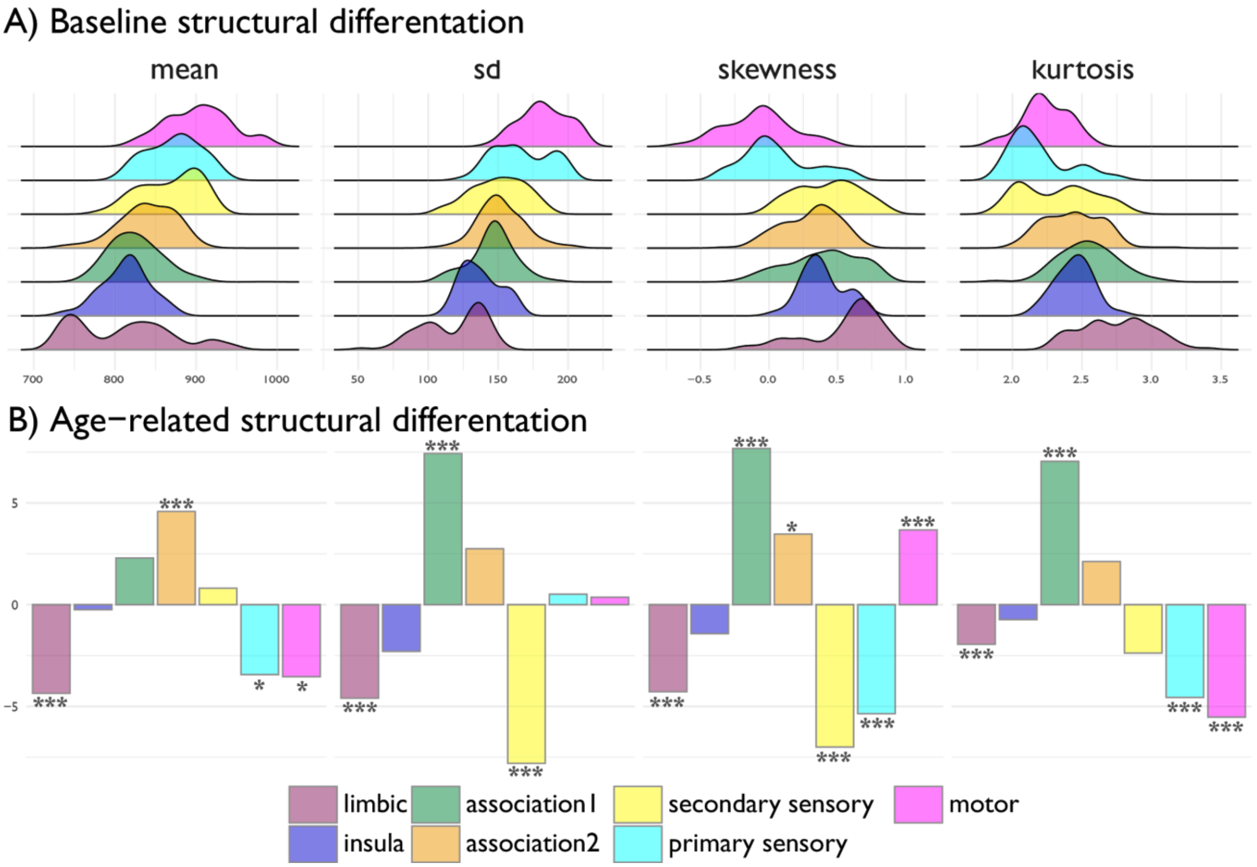


**Supplementary Figure 3:**

Von Economo classification of baseline profiles and age-related t-maps. Panel A shows the distribution of values across each cortical class. Panel B shows the z-scores of relative over or under-representation of parcels showing an age-related effect within each class. Classes showing a significant effect of age related changes are denoted by asterixis (*** = $p < 0.001$, * = $p < 0.05$).

**Supplementary Table 1:**
FDR corrected (Bonferroni) p-Values and Z-scores for age-related changes per level of laminar differentiation

| | P-values | | | | Z-scores | | | |
|---|---|---|---|---|---|---|---|---|
| | mean | sd | skewness | kurtosis | mean | sd | skewness | kurtosis |
| heteromodal | 4.36E-02 | 1.27E-02 | 2.55E-08 | 2.69E-17 | 3.06 | 3.42 | 6.07 | 8.80 |
| idiotypic | 1.53E-07 | 1.00E+00 | 1.45E-04 | 1.76E-17 | -5.78 | -1.91 | -4.49 | -8.85 |
| paralimbic | 1.32E-02 | 4.39E-07 | 7.21E-12 | 1.23E-05 | -3.40 | -5.60 | -7.27 | -4.99 |
| unimodal | 1.90E-04 | 2.85E-01 | 2.43E-02 | 8.38E-01 | 4.43 | 2.45 | 3.24 | 2.03 |

**Supplementary Table 2:**
FDR corrected (Bonferroni) P-Values and Z-scores for age-related changes per Von Economo structural type

| | P-values | | | | Z-scores | | | |
|---|---|---|---|---|---|---|---|---|
| | mean | sd | skewness | kurtosis | mean | sd | skewness | kurtosis |
| association1 | 6.86E-01 | 3.31E-12 | 5.20E-13 | 5.97E-11 | 2.30 | 7.44 | 7.68 | 7.04 |
| association2 | 1.43E-04 | 1.91E-01 | 1.63E-02 | 1.00E+00 | 4.59 | 2.75 | 3.48 | 2.12 |
| insula | 1.00E+00 | 6.97E-01 | 1.00E+00 | 1.00E+00 | -0.25 | -2.29 | -1.42 | -0.73 |
| limbic | 4.14E-04 | 1.39E-04 | 5.98E-04 | 1.00E+00 | -4.36 | -4.59 | -4.28 | -1.95 |
| motor | 1.27E-02 | 1.00E+00 | 7.70E-03 | 1.04E-06 | -3.54 | 0.37 | 3.67 | -5.53 |
| primary sensory | 1.90E-02 | 1.00E+00 | 2.67E-06 | 1.62E-04 | -3.43 | 0.51 | -5.36 | -4.56 |
| secondary sensory | 1.00E+00 | 1.99E-13 | 8.57E-11 | 5.59E-01 | 0.81 | -7.80 | -6.99 | -2.38 |

**Supplementary Table 3:**
Central moments of the profile distribution for age strata at each end of the age distribution corresponding to exemplary regions from figures 1 and 2.

| | **V1** | | **V2** | | **SPL** | | **ACC** | |
|---|---|---|---|---|---|---|---|---|
| | <16 yrs | >24 yrs | <16 yrs | >24 yrs | <16 yrs | >24 yrs | <16 yrs | >24 yrs |
| **Mean** | 920 | 945 | 909 | 926 | 852 | 886 | 824 | 847 |
| **SD** | 158 | 172 | 170 | 186 | 153 | 169 | 129 | 136 |
| **Skewness** | -0.04 | -0.12 | 0.27 | 0.20 | 0.43 | 0.14 | 0.92 | 0.71 |
| **Kurtosis** | 2.01 | 2.01 | 2.02 | 1.96 | 2.22 | 2.07 | 2.91 | 2.45 |

**Correspondence between MT moments**

Comparing spatial topology of both baseline and age-related changes we find that at baseline higher mean MT is spatially coupled with higher variability of MT across cortical depth (mean~SD: r=0.62, p<0.001); skewness and kurtosis exhibit an opposing pattern (mean~skewness: r=-0.27, p<0.001. mean~kurtosis; r=-0.65, p<0.001. SD~skewness: r=-0.64, p<0.001. SD~kurtosis: r=-0.76, p<0.001), with parallel increases in skewness and kurtosis towards heteromodal and paralimbic regions (skewness~kurtosis: r=0.58, p<0.001. **Supplementary Figure 4A**). Although the directionality of these correlations is identical in the t-maps of the age-related change (**Supplementary Figure 4B**), the overall strength is somewhat reduced.

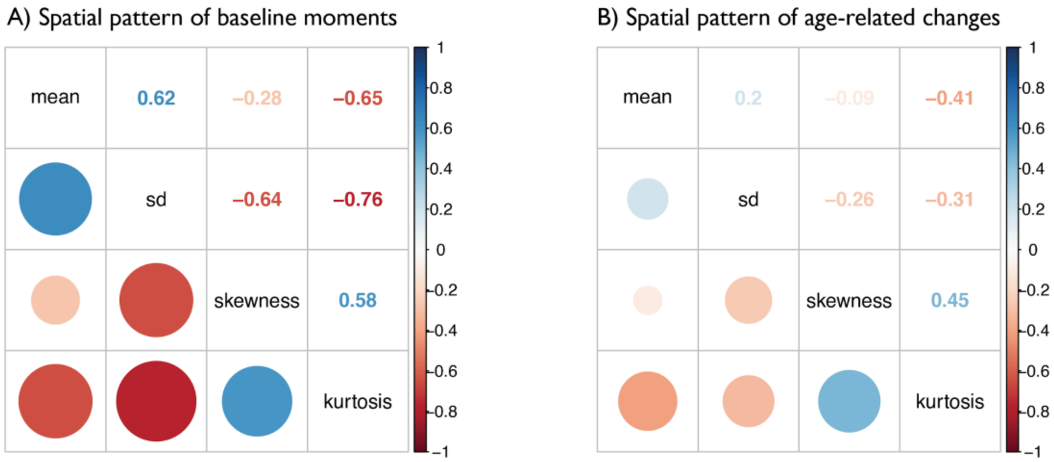


**Supplementary Figure 4:**

Matrices depict the strength of correlations (r values) between *(A)* baseline maps of MT moments and *(B)* t-statistic maps of age-related changes in MT moments.

### Independence of age-related changes from cortical morphology

In light of recent literature showing interactions between cortical morphology and myelin sensitive imaging [9,10], we sought to replicate all our findings while correcting for cortical thickness as well as interface blurring. To assess how regional differences in cortical morphology affect the spatial distribution of MT moments, we regressed out nodal thickness and blurring morphology as such:

$$moment(n) \sim b_0 + b_1 thickness(n) + \mathcal{E}$$

$$moment(n) \sim b_0 + blurring(n) + \mathcal{E}$$

where n represents the node, and recaluted the MT moments as the sum of the residuals. The spatial maps uncorrected MT moments were highly similar to maps corrected for thickness and blurring (all $r>0.99$). For longitudinal assessment, we expanded the existing mixed effects model by including both time-points of data, age and sex as fixed factors and subject as a random factor:

$$moment(n) \sim b_0 + b_1 thickness(n) + b_2 age + b_3 sex + (1|subject) + \mathcal{E}$$

$$moment(n) \sim b_0 + b_1 blurring(n) + b_2 age + b_3 sex + (1|subject) + \mathcal{E}$$

The procedure was repeated for each of the four moments, then we inspected the resultant t-statistics and p values. In addition, we recomputed all effects after regression out the first moment to show that all measures remained independent from any global mean shift. Results show that all age-related changes were independent from cortical thickness (**Supplementary Figure 5**), spatial blurring (**Supplementary Figure 6**) and independent from mean MT (**Supplementary Figure 7**).

#### Cortical thickness

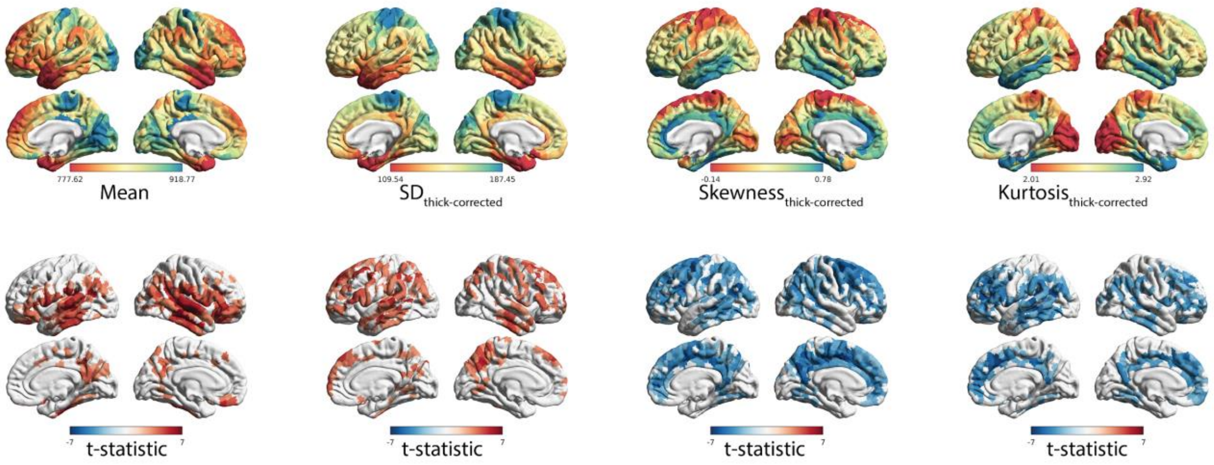


**Supplementary Figure 5:**

Independence of effects controlling for cortical thickness. *(A)* Spatial distribution of MT moments corrected for cortical thickness. *(B) t*-statistics representing the age-related changes in MT moments, with cortical thickness included in the linear mixed effect model as a covariate.

### Interface blurring

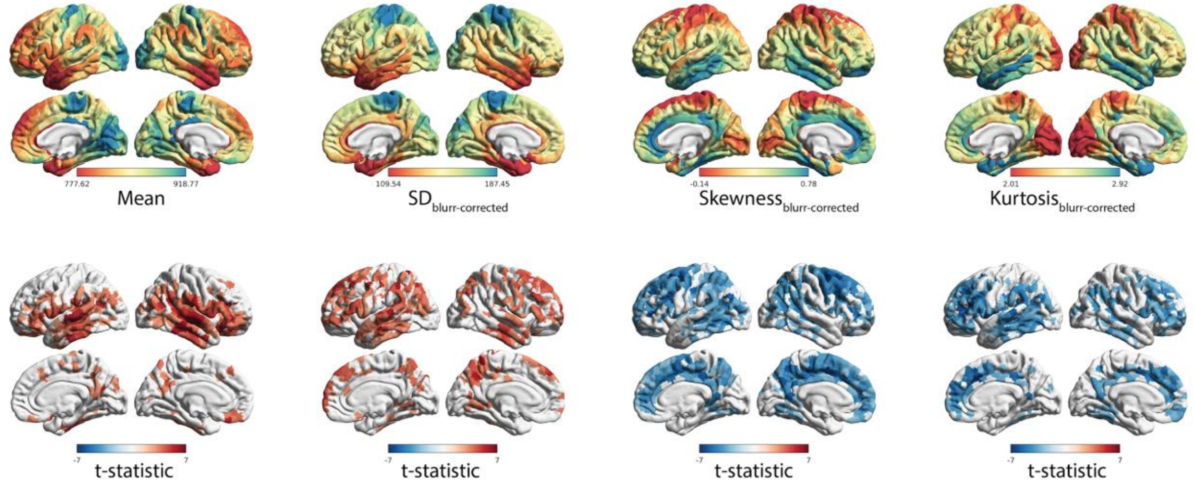


**Supplementary Figure 6:**

Independence of effects controlling for cortical thickness. *(A)* Spatial distribution of MT moments corrected for interface blurring. *(B)* *t*-statistics representing the age-related changes in MT moments, with interface blurring included in the linear mixed effect model as a covariate.

### Global mean shifts

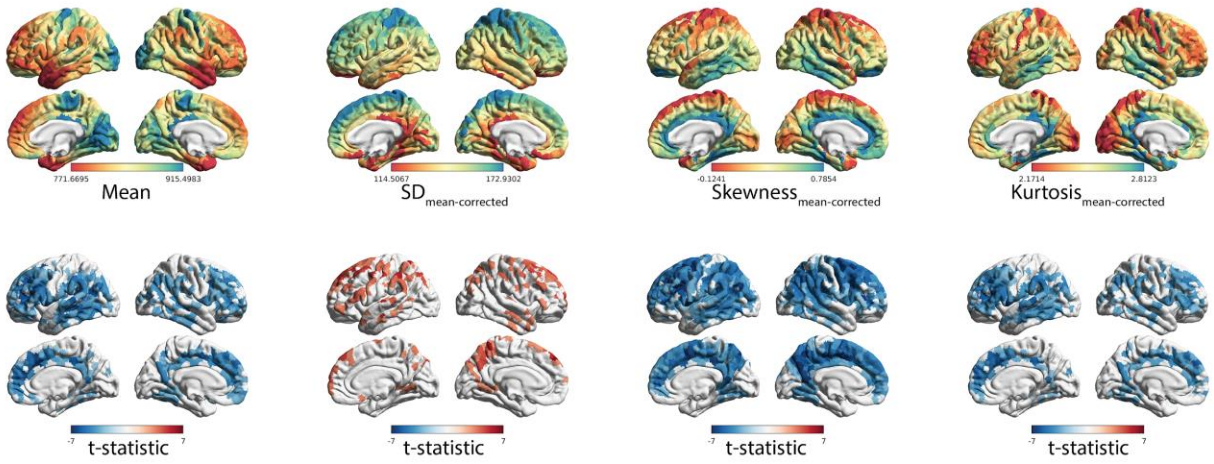


**Supplementary Figure 7:**

Independence of effects controlling for mean MT. *(A)* Spatial distribution of lower MT moments corrected for cortical thickness. *(B)* *t*-statistics representing the age-related changes in MT moments, with mean MT included in the linear mixed effect model as a covariate. Note: mean MT moment results are presented on the left for comparison.

**Baseline gene ontology**

We also assessed the spatial overlap of mean baseline maps with Allen Institute for Brain Sciences (AIBS) gene expression maps [11]. We used Bayesian random effects analysis of AIBS gene expression with spatial topology, as implemented in NeuroSynth gene decoding [12,13], to identify genes significantly associated with each of the maps [11]. We only included genes that passed FDR correction ($P_{FDR} < 0.05$) and conducted enrichment analysis of standard mammalian phenotypes (e.g. the observable morphological, physiological, behavioral and other characteristics of mammalian organisms that are manifested through development and lifespan) in the mammalian phenotype ontology database [14]. Enrichment analyses were conducted using Enrichr [15,16], using a z-score modification of Fisher's exact test. Results from this spatial overlap analysis show no clear or unique ontological profile for any of the genes and when p-values were corrected for FDR across the 4 moments and positive and negative gene-lists, none survived correction (**Supplementary Figure 8**).

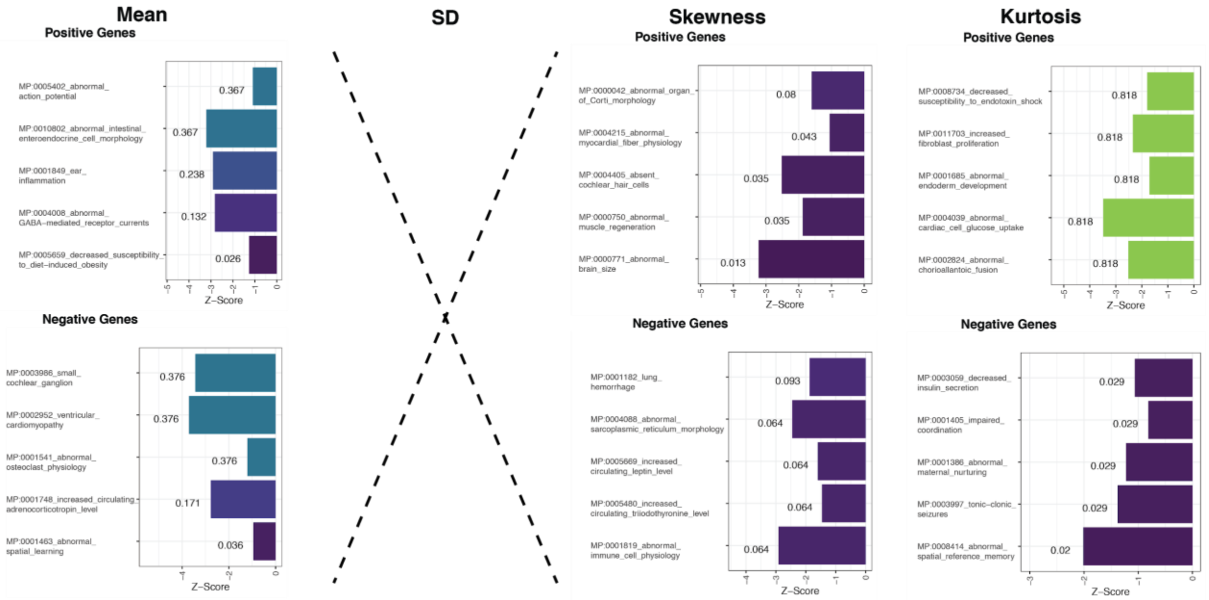


**Supplementary Figure 8:**

Gene ontology analyses from baseline profiles of the four moment of the intensity distribution.

**Deconstructing age-related changes in MPC by cytoarchitectural class**

Age-related changes in MPC showed a scattered but large-scale pattern of age-related changes, mainly in terms of increased microstructure covariance (**Supplementary Figure 9A**). To deconstruct this diffuse pattern, and keeping in line with the deconstruction of results into know cytoarchitectural classes, we also decomposed our developmental MPC matrix by levels of laminar differentiation (Mesulam classes) (**Supplementary Figure 9B**). Mesulam classes did not clearly capture the pattern of age-related changes in MPC. We also sought to decompose the principal gradient by meta-analytic neurosynth decoding. This was done by extracting gradient values using masks based on top neurosynth terms (similar to [17]) and ordering them by mean gradient value to visualize inspect the order of terms across the gradient. We found that previously reported sensory-fugal patterns of the gradient [17,18] are conserved across the age- related change (**Supplementary Figure 9C**).

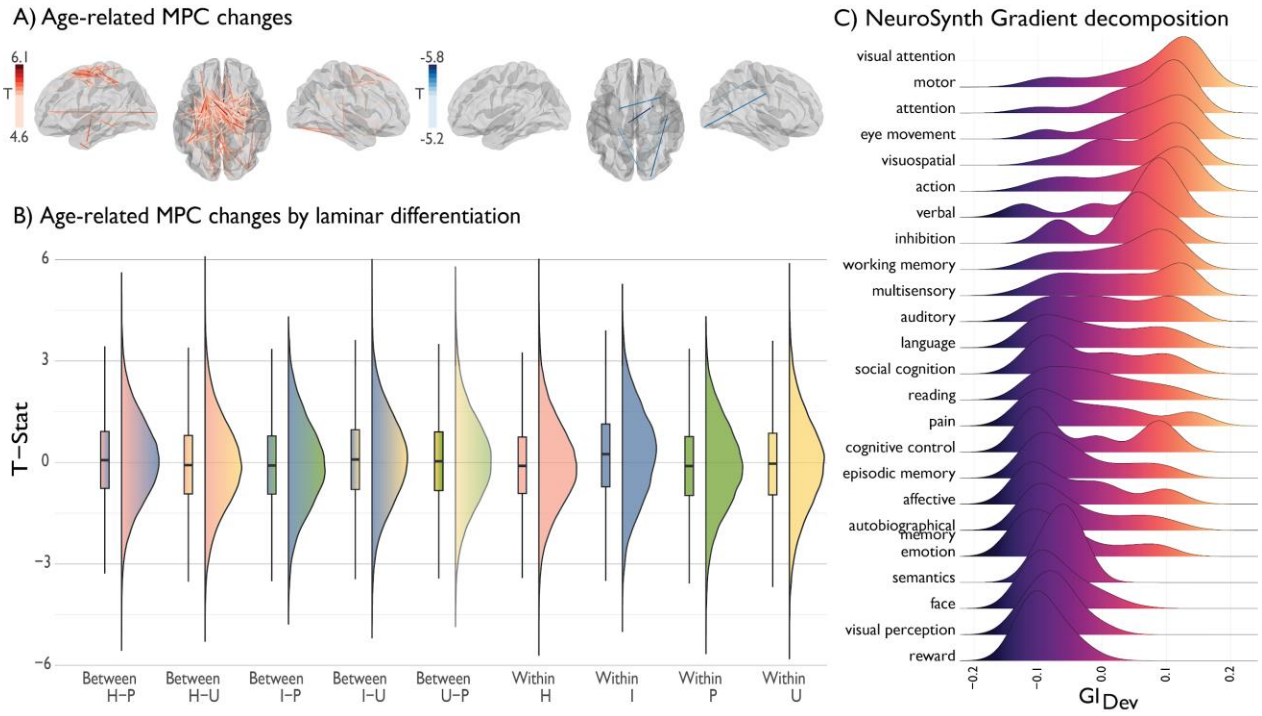


**Supplementary Figure 9:**

A) Age-related changes in MPC stratified by increases (red) and decreases (blue). B) Edge-wise changes in MPC were stratified by level of laminar differentiation. Raincloud plots illustrate the range of t-statistic values within and between different classes. Lines depict the 25%, 50% and 75% quartiles. I=idiotypic. U=unimodal. H=heteromodal. P=paralimbic. C) Ridgeplot depicting distribution of thresholded (z > 2) gradient values across NeuroSynth composite terms, ranked from high to low mean value.

### Increasing bimodality of the principle gradient of microstructural differentiation

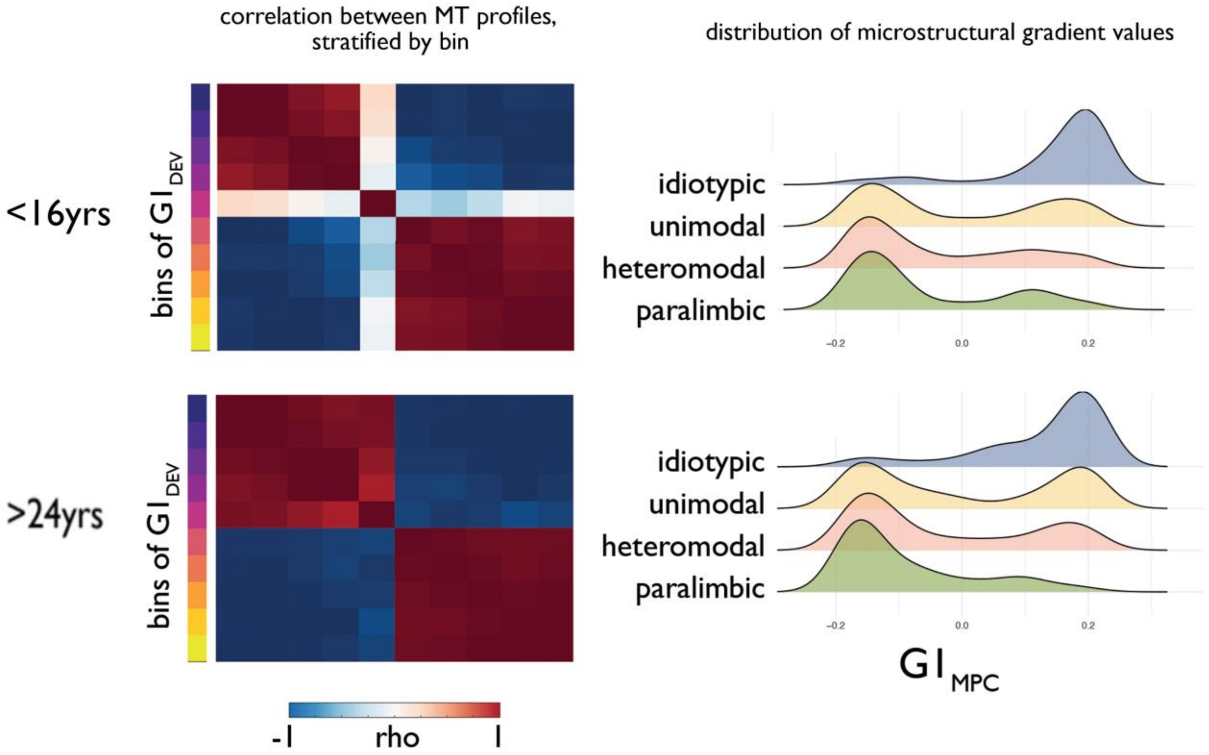


**Supplementary Figure 10:**

***Left*** Matrices showing the Pearson correlation between average MT profiles within ten discrete bins of principal developmental gradient ($G1_{Dev}$) for young (<16years) and old (>24years). ***Right*** Ridge plots of $G1_{MPC}$ by level of laminar differentiation show an increasingly bimodal distribution of $G1_{MPC}$ for association cortices.

**Developmental gradients derived from moment difference**

Absolute difference in MT moment values between each pair of regions was calculated for each participant, and transformed into a node x node matrix. In the same manner as developmental gradient construction, linear mixed effects models estimated the age-related changes of each edge and the resultant t-statistic matrix was subjected to diffusion map embedding. Interestingly, the skewness difference derived principal gradient showed a markedly close and significant resemblance to the principal gradient derived from the age-related change in MPC.

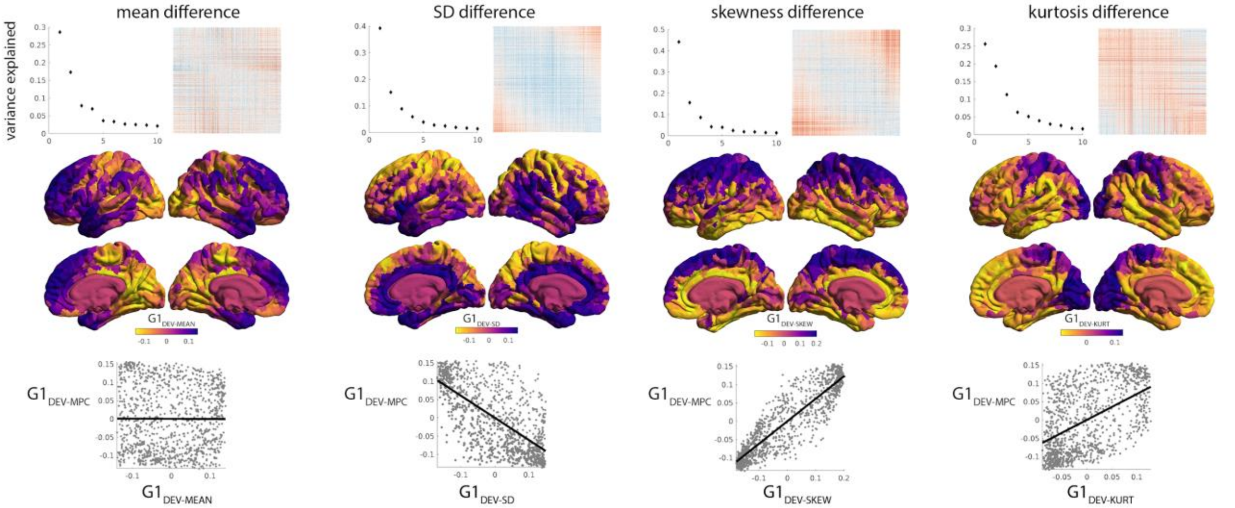


**Supplementary Figure 11:**

Moment-based developmental gradients. Variance explained by diffusion embedding components, t-statistic matrices ordered by the first components, the first components projected onto the cortical surface, scatterplot depicting the similarity of moment-based developmental gradients ($G1_{AGE\text{-}MOMENT}$) to MPC-based developmental gradients ($G1_{AGE\text{-}MPC}$; *Figure 4B*).
